# On the evolutionary origins of equity

**DOI:** 10.1101/052290

**Authors:** Stéphane Debove, Nicolas Baumard, Jean-Baptiste André

## Abstract

Equity, defined as reward according to contribution, is considered a central aspect of human fairness in both philosophical debates and scientific research. Despite large amounts of research on the evolutionary origins of fairness, the evolutionary rationale behind equity is still unknown. Here, we investigate how equity can be understood in the context of the cooperative environment in which humans evolved. We model a population of individuals who cooperate to produce and divide a resource, and choose their cooperative partners based on how they are willing to divide the resource. Agent-based simulations, an analytical model, and extended simulations using neural networks provide converging evidence that equity is the best evolutionary strategy in such an environment: individuals maximize their fitness by dividing benefits in proportion to their own and their partners’ relative contribution. The need to be chosen as a cooperative partner thus creates a selection pressure strong enough to explain the evolution of preferences for equity. We discuss the limitations of our model, the discrepancies between its predictions and empirical data, and how interindividual and intercultural variability fit within this framework.

## 1 Introduction

For centuries, philosophers have emphasized the important role of proportionality in human fairness. In the fourth century BC, Aristotle suggested an “equity formula” for fair distributions (Aristotle, 1999), mathematical equivalent of “reward according to contribution,” whereby the ratios between the outputs O and inputs I of two persons A and B are made equal: 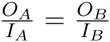. This formula also captures the concept of “merit,” the idea that people who work harder deserve more benefits (Adams, 1963; Konow, 2003; Skitka, 2012).

Psychological research on distributive justice, and on equity theory in particular, has offered extensive empirical support for Aristotle’s claim (Adams, 1963; Homans, 1958; Walster et al., 1973; Mellers, 1982). Equity theory aims to predict the situations in which people will find that they are treated unfairly. A robust finding is that receiving more or less than what one deserves leads to distress and attempts to restore equity by increasing or decreasing one’s contribution (Adams, 1963; Adams and Jacobsen, 1964). People prefer income distributions with strong work-salary correlations, prefer to give more to individuals whose input is more valuable, and favor meritocratic distributions as a whole in both micro-and macro-justice contexts (Baumard et al., 2013).

More recently, experiments with economics games have shown that participants consistently divide the product of cooperative interactions in proportion to each individual’s talent, effort, and the resources invested in the interaction (Cappelen et al., 2010; Frohlich et al., 2004). Meritocratic distributions have been observed across many societies (Marshall et al., 1999), including hunter-gatherer societies (Gurven, 2004; Alvard, 2002; Liénard et al., 2013; Schäfer et al., 2015), and can be detected very early in human development (Kanngiesser et al., 2010; Baumard et al., 2012), suggesting that equity could be a universal and innate pattern in human psychology.

Preferences for equitable outcomes present the same evolutionary problem as preferences for fair outcomes in general: at least in the short term, those preferences are costly. Although people react more to inequitable situations when they are disadvantageous than when they are advantageous, people still feel uncomfortable in unjustified advantageous situations (Austin and Walster, 1974; Fehr and Schmidt, 1999). Experiments even show that people are ready to incur costs and decrease their own payoff in order to achieve more equitable distributions (Dawes et al., 2007). How can natural selection account for the evolution of such costly preferences?

Until now, little attention has been given to this question. There have been many theoretical studies on the evolution of fairness (Nowak et al., 2000; Gale et al., 1995; Page and Nowak, 2002; Barclay and Stoller, 2014; André and Baumard, 2011; Debove et al., 2015a), but all of them are concerned with explaining the evolution of fairness in the ultimatum game, an economic game where the fair division happens to be a division into two equal halves (Güth et al., 1982; Camerer, 2003). However, equal divisions are just a special case of the more general category of equitable divisions: that is, divisions proportional to contributions. As emphasized by equity theory, unequal divisions can be judged fair when they respect the partners’ investment, talents, commitment, etc. In brief, although many models can explain the evolution of preferences for *equal* divisions, none of them is able to explain the evolution of preferences for *proportional* divisions. Here we aim to understand whether natural selection can lead to such proportional divisions of resources (including the particular case of equal divisions), in a scenario where partners can make differing contributions to a cooperative undertaking.

Partner choice has had an important role in the evolution of cooperation, as evidenced by both theoretical (Aktipis, 2004; Nesse, 2007; Aktipis, 2011; McNamara et al., 2008; Barclay, 2011) and empirical studies (Barclay, 2004; Barclay and Willer, 2007; Sylwester and Roberts, 2013, and see Barclay, 2013 for a review in humans). When people are in competition to be chosen as cooperative partners, experiments show that they increase their level of cooperation because they have a direct interest in doing so (Barclay, 2004, 2006). Partner choice also has interesting consequences for the evolution of fairness. It leads to equal divisions of resources in theoretical and empirical settings (André and Baumard, 2011; Debove et al., 2015b,a), because when individuals can choose whom to cooperate with then they are better off refusing divisions that do not compensate their opportunity costs. These results suggest the way through which partner choice could also explain the evolution of divisions proportional to contributions: if greater contributors have larger opportunity costs, they will choose partners who give them something at least equal to these opportunity costs. Nonetheless, this hypothesis has never been studied formally.

To summarize, preferences for equity are robust and widespread in humans, but we currently lack an evolutionary explanation for their costly existence. Here, we aim to put the partner choice mechanism to the test to see if it can explain such preferences. We develop models in which individuals put effort into the production of a collective good, and differ with regard to both the amount of effort they are willing to put in and the efficiency of their contribution to the production of the good. To determine the evolutionary stable sharing strategy in this environment, we first analyzed an evolutionary model using agent-based simulations. We then developed a simple analytical model to better understand the simulations, and tested the robustness of our results by performing simulations with evolving neural networks as more realistic decision-making devices. The results provide converging support for the conclusion that when individuals can choose whom to cooperate with, equity emerges as the best strategy, and the offers that maximize fitness are those that are proportional to the individual’s relative contribution to the production of the good.

## 2 Methods

We develop three complementary sets of simulations and an analytical model. For clarity, we present the first set of simulations in details before explaining how the other sets differ. Source code for all simulations is available online.

### 2.1 Simulations Set 1: two productivities

#### 2.1.1 Individuals

We consider a population of *n* individuals who will be given multiple opportunities to cooperate and produce resources during their life. Cooperation only takes place in dyadic interactions. We assume individuals are characterized by a “productivity”, such that some individuals can produce more resources than others when they cooperate. Individuals can be of one of two productivities: low-productivity individuals can produce *a* resources when they cooperate, while high-productivity individuals can produce *b* resources *(b > a)*. This productivity is constant across the entire life of an individual but is not heritable: at birth, each individual is randomly attributed a level of productivity that is independent of his parent’s. This condition is necessary so that there is always a diversity of productivities in the population at each generation.

To decide with whom they will cooperate and how to divide resources, we assume that each individual is characterized by eight genetic variables: four *r*_*ij*_ and four MAR_*ij*_ variables, with *i* and *j* ∈ {*HP, LP*}, denoting an individual’s productivity (HP = High-Productivity, LP = Low-Productivity). *r*_*ij*_ is the fraction of resources (between 0 and 1) that an individual of productivity *i* will give to an individual of productivity *j*. We call the *r*_*ij*_ variables the “reward” variables. MAR_*ij*_ is the minimum acceptable reward, the minimum fraction of resource that an individual of productivity *i* is ready to accept from an individual of productivity *j*.

#### 2.1.2 Social life

Only two types of events can happen at any given time in our model: the encounter of two solitary individuals, or the split of two cooperating individuals. We model time continuously. At each loop of the model, we (i) determine the time period until the next event (ii) determine whether this event is an encounter or a split, and (iii) execute the corresponding actions for each event, described below. This process is repeated until time has exceeded a constant *L*, which corresponds to the end of the life of all individuals (see section “reproduction” below).

After any event occurring at time *t* (or after the birth of individuals at t=0), the time period until the next event is drawn in an exponential distribution of parameter

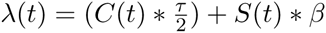

with *C(t)* the number of cooperating individuals at time *t, S(t)* the number of solitary individuals at time *t, β* a constant encounter rate and *τ* a constant split rate.

The probability *p(t)* that this event is an encounter is then given by

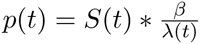

Conversely, 1 *— p(t)* is the probability that this event is a split.

Depending on whether the event is an encounter or a split, two scenarios unfold:

1/ If the event is an encounter, two solitary individuals are randomly drawn from the population and offered an opportunity to cooperate to produce resources. To this end, one of the two individuals is randomly selected to unilaterally decide how to divide the resources through her *r*_*ij*_ reward variable. We call this individual the “partner”. However, before cooperation effectively starts, the partner must be accepted by the second individual. We call the second individual the “decision maker”. The decision maker makes her decision based on her partner’s reputation. For simplicity, we do not model the formation of this reputation. We simply assume that the decision maker knows her partner’s reward value *r*_*ij*_ For instance, a HP partner *A* has a reputation of *r*_*A*_*H P L P*__ with a LP decision maker *B*. The LP decision maker will then compare the value of *r*_*A*_*H P L P*__ to her own *MAR*_*B*_*L P H P*__, and if *r*_*A*_*H P L P*__ ≥ MAR_*B*_*L P H P*__, the partner will be accepted and cooperation will start. From this point on until the interaction stops, the two individuals produce, at each unit of time, an amount of resources that is equal to the sum of their respective productivities, from which the decision maker receives a fraction *r*_*A*_*H P L P*__.Conversely, if the partner’s reputation is not good enough for the decision maker (*r*_*A*_*H P L P*__ < MAR_*B*_*L P H P*__), the two individuals do not cooperate together and go back to the pool of solitary individuals without receiving any resources.

2/ If the event is a split, a pair of cooperating individuals is randomly chosen to split, and the two individuals go back to the pool of solitary individuals.

#### 2.1.3 The cost of partner choice.

The cost of partner choice is implicit in our model. It is a consequence of the time it takes to find a partner. Hence, the cost and benefit of being choosy are not controlled by explicit parameters, but by two parameters that characterize the “fluidity” of the social market: the “encounter rate” *β*, and the “split rate” *τ*. When 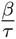 is large, interactions last a long time (low split rate *τ*) but finding a novel partner is fast (high encounter rate *β*), and individuals thus should be picky about which partners they accept. This is a situation where partner choice is not costly. On the contrary, when 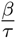 is low, interactions are brief but finding a novel partner takes time, and individuals should thus accept almost any partner. Partner choice is then costly.

#### 2.1.4 Reproduction

We model a Wright-Fisher population with non-overlapping generations: when the lifespan *L* has been reached, all individuals reproduce and die at the same time. The number of offsprings produced by a focal individual is given by:

> offsprings = round 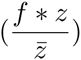

with *z* the focal individual’s amount of resources accumulated throughout her life, 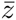 the average amount of resources accumulated in the population, and *f* a constant multiplication factor. Offsprings receive the four *r*_*ij*_ and four MAR_*ij*_ traits from their parents, with a probablity *m* of mutation on each trait. Mutations are drawn from a normal distribution centered around the trait value with standard deviation *d*, and constrained in the interval [0,1]. After mutations take place, *n* individuals are randomly drawn from the pool of offsprings to constitute the population for the next generation.

Table 1 summarizes the model’s parameters. To obtain the results presented below, we initialize all simulations with a population of stingy and undemanding individuals, who do not share when they play the role of partner and accept any partner when they play the role of decision maker (*r*_*ij*_ = 0, MAR_*ij*_ = 0). We then test our hypothesis that partner choice can lead to equitable divisions by observing how rewards and MARs evolve across generations, in two conditions: when partner choice is costly (low 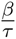), and when partner choice is not costly (large 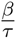). In particular, we will observe the rewards given by LP individuals to HP individuals at the equilibrium when partner choice is not costly, to detect whether they show the same pattern of proportionality between contribution and reward than the one observed in the empirical human data.

**Table 1:** Parameters of the model, and values used to obtain the figures presented in the main text. Deviations from these values do not change the core results.

| Parameter name | Description | Value used to obtain reported results |
| --- | --- | --- |
| n | number of individuals | 500 |
| a | productivity of low-productivity individuals | 1 |
| b | productivity of high-productivity individuals | 2 |
| r | reward, fraction of resources that an individual agrees to give to another | evolving (starts at 0) |
| MAR | minimum accepted reward, minimum fraction of resource that an individual is ready to accept | evolving (starts at 0) |
| $\beta$ | encounter rate | from 0.0001 to 1 |
| $\tau$ | split rate | 0.01 |
| L | lifespan | 500 |
| m | mutation rate | 0.002 |
| d | mutation standard deviation | 0.02 |

### 2.2 Analytical model.

We develop an analytical model that incorporates all of the features of the simulations presented above, but with one simplification: we assume that the total number of interactions accepted per unit of time is the same for each individual. With this assumption, rejecting an opportunity to cooperate does not compromise the chances of cooperating later, but on the contrary grants new opportunities. This situation is analogous to the condition where 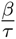 tends towards infinity in the simulations: social opportunities are plentiful at the scale of the length of interactions. The analysis of this model is presented in details in SM section B.

### 2.3 Simulations Set 2: a continuum of productivities

Introducing a continuum of productivities is necessary to get closer to biological reality. Rather than having only two productivities *a* and *b* in our population, we assume in Simulations Set 2 that the productivity of an individual at birth is sampled from a uniform distribution between *a* and *b*. In this situation, individuals never interact with a partner of the exact same productivity. This constitutes a challenge for modeling in that individuals would need to be equipped with an infinity of *r*_*ij*_ and MAR_*ij*_ traits to react to the infinity of possible contributions by their partner (Gavrilets and Scheiner, 1993).

To solve this problem, we do not characterize anymore individuals with *r*_*ij*_ and MAR_*ij*_traits, but instead endow them with two three-layer feedforward neural networks (one network to produce the rewards, and another one to produce the MARs). Both neural networks have the same structure: two input neurons, five hidden neurons, and a single output neuron. The first neural network is used when playing the role of partner: it senses an individual’s own productivity and that of her decision maker, and produces the reward as output. The second network is used when playing the role of decision maker: it senses an individual’s own productivity and that of her partner, and produces the MAR as output. Each network has its own set of synaptic weights (see Fig. 3A and SM section A.2), that are transmitted genetically. Because evolution now operates on these weights, and not on rewards or MARs directly, individuals can now evolve a reaction norm. They can evolve a function that produces outputs even from inputs they have never encountered before (i.e., individuals of new productivities). This property of neural networks is important in our case, because equity is precisely a relationship between two quantities, contribution and reward. Seeing whether natural selection will be able to recreate the same relationship of proportionality between contribution and reward using simple neural networks is thus of great interest. All other methodological details for Simulations Set 2 are the same as in Simulations Set 1.

**Figure 3:**
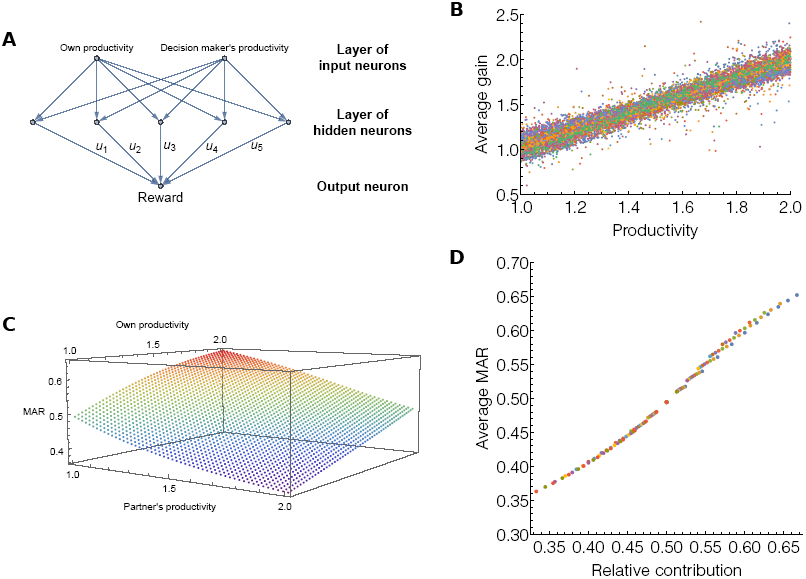
Evolution of equitable rewards made by neural networks working on a continuum of productivities. A: Schematic representation of the neural networks that make rewards. Networks take each individual’s productivity as inputs and produce the reward as output. The u’s represent synaptic weights on which evolution takes place. B: 15,000 individuals and their lifelong average gain plotted against their productivity. C: Average MARs produced by the neural networks of 15,000 individuals after 8,000 generations, for different values of the input neurons. The more an individual produces and the less the partner produces, the larger the individual’s MAR. D: Average MARs produced by 15,000 neural networks plotted against the relative contribution of the bearer of the network.

### 2.4 Simulations Set 3

As a final test of the robustness of our model, we test whether natural selection also favors divisions proportional to contributions when contribution is measured in terms of time invested into cooperation (instead of productivity). We present the details of these simulations and its results in SM section A.1.

## 3 Results

We first present the results for Simulations Set 1. Parameter values used to obtain the figures are summarized in Table 1. Reasonable deviations from these values do not alter the results. Moreover, analytical results confirm the results of Simulation Set 1 (see SM section B).

We present the case where high-productivity individuals are able to produce twice as much resources as low-productivity individuals (*a* = 1, *b* = 2). Figure 1 shows the evolution of rewards *r* accepted by decision makers across generations. Rewards increase in all possible combinations of productivities, when partner choice is not costly (circle markers). If we focus on rewards accepted by high-productivity decision makers with low-productivity partners (Fig 1, upper-right panel), simulations show that at the evolutionary equilibrium, low-productivity partners have to give exactly 66% of the total resource produced to their high-productivity decision makers. This reward is exactly proportional to the relative contribution of each individual, as high-productivity individuals produce 66% of the total shared resource when *a* = 1 and *b* = 2. Similarly, high-productivity partners give only 33% to low-productivity decision makers, a reward which low-productivity decision makers accept, as it corresponds to their relative contribution (Fig 1, lower-left panel, circle markers). Finally, both high-productivity and low-productivity individuals give each other exactly 50% of the total resource when they meet as a pair, reflecting the fact that proportionality means equal division when contributions are equal (Fig 1, upper-left and lower-right panels). This pattern of divisions is confirmed by the analytical model (dashed lines in Fig 1, and see SM section B), and divisions proportional to contribution also evolve when contribution is measured in terms of time invested into cooperation instead of productivity (see SM section A.1).

**Figure 1:**
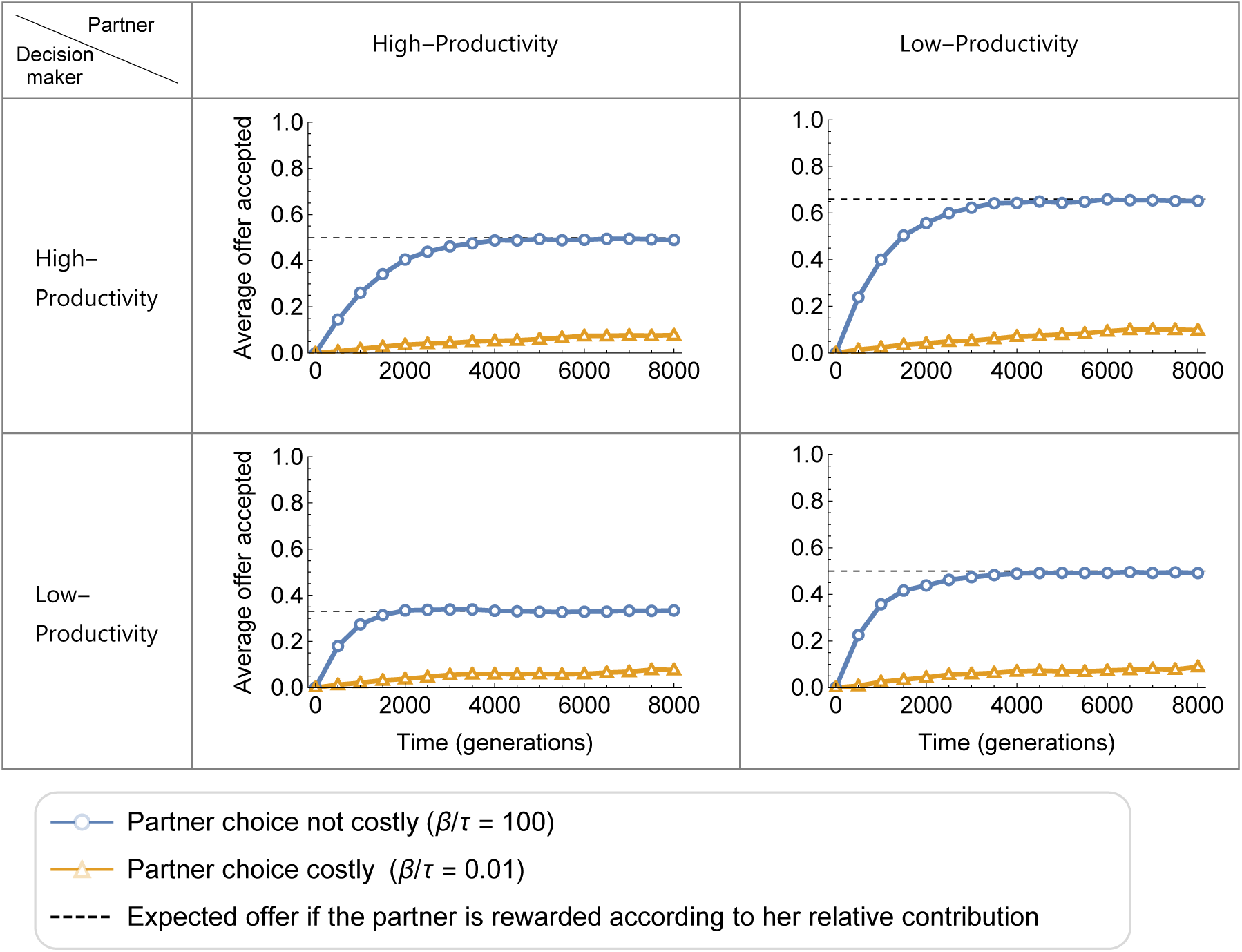
Evolution of the average rewards accepted in cooperative interactions according to the productivity of the decision maker and the partner. High-productivity individuals produce twice as much resources as low-productivity individuals. When partner choice is not costly, rewards evolve to match the decision maker’s relative contribution. Dashed lines represent the expected reward in the analytical model. The evolution of MARs is visually undistinguishable from the evolution of rewards and thus not represented.

By comparing simulations with a low and a high 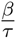 ratio, Figure 1 also emphasizes the critical importance of partner choice for proportional rewards to evolve. When we decrease the 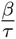 ratio, individuals spend more time looking for new partners and thus the cost of changing partners is increased. In this situation, rewards remain very low over generations and never rise towards proportionality, regardless of differences in productivity (Fig 1, triangle markers). For instance, even if low-productivity partners produce less than half of the resources when they cooperate with high-productivity decision makers, they keep most of the resources for themselves when partner choice is costly. Figure 2 shows the distribution of rewards given by low-productivity individuals to high-productivity individuals at the end of an 8,000-generation simulation, for different values of the 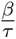 ratio. Proportional rewards of 66% can only evolve when 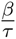 is large, showing again that without partner choice, proportionality cannot evolve.

**Figure 2:**
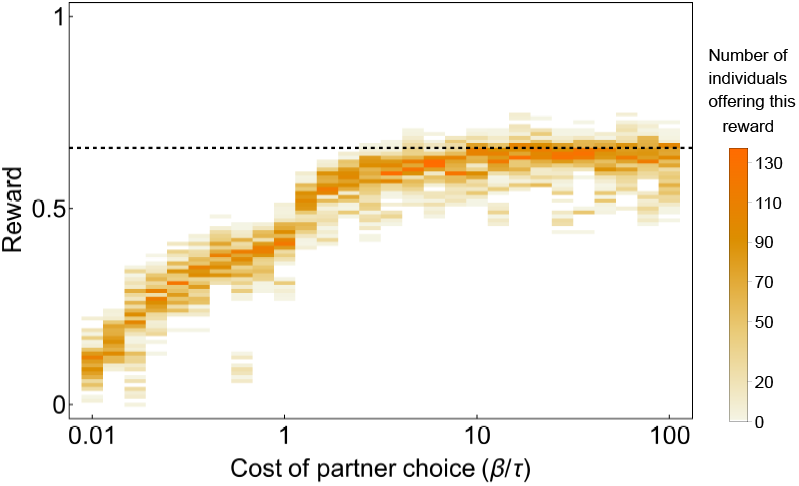
Distribution of rewards offered by low-productivity individuals to high-productivity individuals in the last generation of an 8,000-generation simulation, for different levels of partner choice cost (higher values of 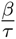 represent lower costs). High-productivity individuals’ relative contribution compared to low-productivity individuals is 0.66, so the dashed line represents the expected equitable distribution. This distribution can only be reached when partner choice is not costly (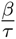 is high).

The results of Simulation Set 2 confirm this pattern. With a continuum of productivities in the population (between 1 and 2), rewards still respect proportionality at the evolutionary equilibrium. Each individual who enters an interaction is rewarded with an amount of resources exactly equal to her productivity (Fig 3B). As explained in the methods section, neural networks have two inputs: an individual’s own contribution and her partner’s (or decision maker’s) contribution. It is thus possible to represent the output of a network on a 3D plot, shown in Fig 3C. To plot this figure, we extracted the synaptic weights of the neural networks producing MARs for 15,000 individuals, at the last generation of 30 different simulation runs. We averaged the value of the networks’ outputs over those 15,000 individuals. Fig 3C shows that the networks evolved to produce MARs that are proportional to their bearer’s relative contribution (Fig 3C and D, and see SM section C.2). The higher the decision maker’s productivity, and the lower the partner’s productivity, the more demanding the decision maker becomes.

## 4 Discussion

We modelled a population of individuals choosing each other for cooperation. When different contributions to cooperation are made, resource divisions proportional to contributions evolve. Individuals producing more resources or investing more time into cooperation receive more resources than individuals producing or investing less. Asking for divisions that match one’s own contribution, and proposing such divisions to others, constitutes the best strategy when partner choice is possible. In other terms, a preference for equity maximizes fitness in an environment where individuals can choose their cooperative partners.

It is important to note that our results cannot be summarized as “a preference for equity helps individuals to be chosen as a partner” or “a preference for equity helps avoid interactions with selfish partners.” This is only half of the story. If the point were only to be chosen as a partner, the best strategy would be to be as generous as possible, an outcome which is sometimes observed in models inspired by competitive altruism theories (Roberts, 1998). The point here is rather to be chosen as a partner while at the same time avoiding exploitation by being over-generous. Our model clearly shows that the best strategy to solve this problem is to give proportionally to the other’s contribution—not less, but also not more. Equity is the result of a trade-off between two evolutionary pressures which work in opposite directions: the pressure to keep being chosen, but also the pressure to choose wisely.

This last point is better understood by looking at the precise mechanism through which proportionality evolves. The key factor determining divisions of resources at the evolutionary equilibrium are the opportunity costs of each individual. Opportunity costs represent the benefits an individual renounces to when she makes a choice. From an evolutionary point of view, it is trivial that an individual will want to make the best choices possible to minimize her opportunity costs. Hence, the best strategy to keep being chosen as a cooperative partner is to compensate others’ opportunity costs: when individual A agrees to interact with individual B, individual B should give A something equal to A’s opportunity costs at the time of making the decision (and vice versa). This is exactly why high-productivity individuals get more in our model: high-productivity individuals have larger opportunity costs than low-productivity individuals. Suppose that low-productivity individuals produce 1 unit of a resource whereas high-productivity individuals produce 2. High-productivity individuals thus have the possibility to produce 4 resources when they interact with other high-productivity individuals, leaving them with 2 resources on average (see exactly why in SM section C.1). 2 resources is thus the opportunity cost of high-productivity individuals when they agree to cooperate with low-productivity individuals. Thus, if low-productivity individuals want to be good partners, they will have to compensate high-productivity individuals’ opportunity costs and give them exactly 2 resources (out of 3 produced), which will result in a proportional offer of 66%. But low-productivity individuals should not give more neither, because they also have access to interactions in which they could gain 1 unit on average (when they cooperate with other low-productivity individuals). In other words, low-productivity individuals have opportunity costs of 1, and should thus not accept divisions leaving them with less than 1. Our current model and previous papers on the subject (André and Baumard, 2011; Debove et al., 2015b,a) push forward the idea that the sense of fairness is a psychological mechanism evolved to compensate others’ opportunity costs and minimize one’s own opportunity costs. This characterization only comes from models investigating fairness in distributive situations though, so it would be interesting to see if it holds in more diverse, non-distributive situations.

Our model has several limitations, which need to be acknowledged. First, while we suppose that individuals choose each other based on their reputation, we do not explicitly model the formation of this reputation. Individuals automatically know the reputation of others and this reputation is reliable. It could be interesting to relax this assumption, especially because reputation formation (through communication for instance) might be an important point that distinguishes humans from non-human primates. Second, the population we model does not match the hunter-gatherer population in the sense that it is not structured. This is important because a structure, such as camps or family units, could potentially affect opportunities to choose partners. Finally, it might be interesting to model the evolution of fairness in a wider range of cooperative interactions than we have considered here (outside distributive situations for instance). All of these assumptions should be relaxed in future studies.

Partner choice is not the only evolutionary mechanism postulated to lead to the evolution of fairness in the literature. Some authors have argued that fairness could be explained by empathy (Page and Nowak, 2002), spite (Huck and Oechssler, 1999; Barclay and Stoller, 2014; Forber and Smead, 2014), “noisy” processes such as drift or learning mistakes (Gale et al., 1995; Rand et al., 2013), the existence of a spatial population structure (Page et al., 2000; Killingback and Studer, 2001), or alternating offers (Rubinstein, 1982; Hoel, 1987). But as we explained in the introduction, all of these models equate fairness with equality, and it is thus unknown whether they can explain a more general case. Testing whether those models pass the “equity test” will be an excellent way to compare and decide between these models, a necessary undertaking that has been largely neglected. The extensive literature on “bargaining” in economics (Binmore, 1986; Binmore, 1998; Alexander, 2000) was also more focused on the case in which players are in a symmetric position, and usually did not investigate proportional bargaining solutions. An exception is the work by Kalai (1977) (although Binmore, 2005 also mentions the problem p. 31), who shows that individuals will compromise in different bargaining situations so as to keep their proportions of utility gains fixed. But, as Kalai recognizes it himself (P11), “a more difficult problem is to find what these proportions should be”. This is precisely where we make a contribution: we show that when individuals evolve in biological markets, these proportions are automatically determined by the other encounters individuals can make. In other words, one could rephrase our model as showing that individuals can bargain based on their outside options (or opportunity costs), but contrarily to what has been done before, we do not fix exogenously those outside options. Rather, outside options emerge endogenously from all the encounters individuals can make in the population.

Talking about bargaining theory suggests alternative interpretations of our model. It might be argued that human fairness is the result of bargaining at the proximal level, the result of rational cognitive processes. We argue instead that the “bargaining” already took place at the ultimate level by means of natural selection, and that the result of this bargaining is the existence of a genuine sense of fairness which “automatically” makes humans prefer equitable strategies. This hypothesis does not exclude the possibility that humans are also capable of consciously bargaining based on their opportunity costs, but this behavior would not be the product of an evolved sense of fairness. While our model bears a great resemblance to historical market models (Osborne and Rubinstein, 1990) and other models in economics in which fair outcomes have sometimes been observed (Rubinstein, 1982; Binmore, 2005), we emphasize that the markets we model are ultimate biological markets (Noë and Hammerstein, 1994; Noë et al., 2001). This is not just an empty terminological variation: locating markets at the ultimate level has important implications for our understanding of the psychological mechanisms underlying fairness. Among other things, it allows us to understand why fairness does not seem to be based on self-interest at the psychological level even if fairness evolved for self-interested reasons (Baumard et al., 2013; Trivers, 1971).

Another alternative interpretation of our model remains. One could agree that fairness judgments are based on simple automatic rules rather than complex conscious calculations, but argue that those rules could have evolved culturally rather than biologically. This is not an issue that can be settled theoretically, as the same models can always be interpreted as instances of biological or cultural evolution. To date, we definitely lack empirical data to answer this question with certainty, but the idea of a biologically evolved sense of fairness is not made absurd by the existing data. As early as the age of 12 months, children react to inequity (Schmidt and Sommerville, 2011; Geraci and Surian, 2011; Sloane et al., 2012), equity has been identified in many cultures around the word (Marshall et al., 1999; Gurven, 2004), and children reject conventional rules when they violate principles of fairness (Turiel, 2002). We do not take experiments on inequity aversion in non-human primates as evidence for a biologically evolved sense of fairness, as the negative reactions to inequity observed so far can still be interpreted in more parsimonious ways (see Bräuer and Hanus (2012) for a review and Amici et al. (2014) for methodological issues). Nonetheless, those experiments remind us that many researchers expect that prosocial behaviors traditionally associated with the existence of human institutions, religions, or cultural artefacts can also evolve biologically. In fact, Robert Trivers himself recognized that the most important implication of his seminal paper on the evolution of reciprocity (Trivers, 1971) was that “it laid the foundation for understanding how a sense of justice evolved” (Trivers, 2006).

The existence of intercultural and interindividual variations in fairness judgements (?Cappelen et al., 2010; Schäfer et al., 2015) is sometimes taken as evidence against their biological origin. This criticism is generally ill-founded, as evolutionary explanations have no particular difficulty accommodating variation (Barkow et al., 1992). In the case of fairness, it is important to remember that what our model predicts is not the evolution of a fixed judgement but the evolution of an algorithm, an information-processing mechanism (Barkow et al., 1992). This is particularly evident in our extended simulations where the evolving unit is a neural network, precisely a special type of algorithm. This algorithm works on inputs (contributions) to produce outputs (divisions of resources), and here lies an important source of variability, because inputs can vary across cultures and individuals while the algorithm remains the same. For instance, measurements of contributions are affected by beliefs (“How long do I think it takes to harvest this quantity of food?”). If contribution was the only input in our model, in real-life more parameters can affect the algorithm’s inputs, such as general knowledge (“Is this person not productive because she is sick?”) or individual interpretations of the situation (“Are we engaged in a communal interaction? A joint venture? A market exchange?”). This last point could explain why even in carefully controlled environments, where there is little ambiguity about the source of inequalities, there is still heterogeneity in fair behaviors, with some people behaving as egalitarians, others as meritocrats, and others still as libertarians (Cappelen et al., 2007, 2010).

In fact, while interindividual and intercultural variations have crystallized the debate, intra-individual variation can also be observed even in Western countries. In some situations we behave as meritocrats, requiring pay for each additional hour of presence at work (Adams, 1963; Adams and Jacobsen, 1964), whereas the next day on a camping trip with strangers we behave more like egalitarians, without constant monitoring and bookkeeping of our contributions and those of others (Cohen, 2009). Neither our brain (the algorithm) nor our culture has changed in the meantime. What has changed is the way we interpret the situation (part of the input to the algorithm). This idea needs to be developed more formally, and we do not suggest that it is the only way to explain variation, but it may constitute a fruitful avenue of research.

Another interesting question is the prevalence of equity in traditional societies. We have mentioned anthropological records of distributions according to effort (Gurven, 2004; Kaplan and Gurven, 2005), but it is also well known that hunter-gatherers transfer meat in a way that not does not seem to respect equity. This type of interaction has been called “generalized reciprocity” by Sahlins (1972) and also seems to match Fiske (1992)’s notion of a “communal sharing” system. There are at least two mutually compatible ways to reconcile this observation with the predictions of our model. The first is to recognize that equity can be limited by other factors, for instance diminishing returns to consumption (Nettle et al., 2011). People could stop caring about equity when they become satiated or when they receive little additional value from consuming one more unit of benefits. The second is to consider that even in generalized reciprocity good hunters are rewarded with more benefits, but those benefits are delayed. This hypothesis has received support recently from findings showing that generous hunters and hard workers are central in the social networks of small-scale societies (Lyle and Smith, 2014; Bird and Power, 2015). In this last perspective, our model should not be taken at face value as predicting the evolution of strict equity with immediate input/output matching, but more generally as input/output matching over a long time and across different cooperative activities (“generalized equity”).

We conclude by noting that proportionality is important in distributive justice but is also a cornerstone of institutional justice, wherein offenders are punished in proportion to the severity of their crimes (Hoebel, 1954; Robinson and Kurzban, 2007). It is also central to the morality of many religions, in which rewards and punishments are made proportional to good and bad deeds by supernatural entities or forces (Baumard and Boyer, 2013). Although this is only speculation at present, our results may thus also explain why historically recent cultural domains such as penal justice and moral religions insist on the principle of proportionality: retributive punishment and supernatural justice may reflect our evolved desire for proportionality.

## Supplementary Material

### A Simulation procedures

#### A.1 Simulations Set 3: contribution through time invested

##### A.1.1 Methods

Having a higher productivity is only one way to contribute more to a cooperative interaction. Another natural way is to spend more time to amass resources. To test the robustness of our partner choice mechanism, we thus created a third set of simulations in which there are no more differences of productivity between individuals, but one of the two individuals in a cooperating dyad has to invest *m* times more time than her partner. We thus model the possibility that there is a cooperative role more time-consuming than the other. In practice, we model this by randomly attributing a “high investment of time” role to the partner or the decision maker when an encounter takes place. The decision maker then decides whether or not she wants to cooperate with her partner based on her partner’s reputation for a given level of investment into cooperation. Each individual is thus characterized by 4 genetic variables, two *r*_*kl*_ and two MAR_*kl*_, with *k* and *l* ∈ {*H, L*}, denoting an individual’s time investment (H = High, L = Low). If the partner is accepted, individuals share a constant resource of size 1 at each unit of time, and the end of the interaction is determined in the same way than in Simulations Set 1, through a constant split rate *τ*. When a split happens though, the individual who needs to invest more time is prevented to encounter new individuals for a length of time equal to (*m* − 1) * (the length of the interaction). Because this individual is prevented to encounter other individuals during this period, one can interpret this period as a period in which this individual is still investing time into the previous interaction.

All other methodological details for Simulations Set 3 are the same as in Simulations Set 1. In particular, we start from a population of individuals giving zero reward even when they invest less time into cooperation, and observe what will be the relationship between contribution (time invested) and rewards at the evolutionary equilibrium.

##### A.1.2 Results

Simulations Set 3 show that proportional rewards also evolve when individuals differ not by their productivity but by the time they invest in cooperation (Fig 4). Setting *m =* 2, one individual of the pair has to invest twice as much time as the other. When the decision maker invests twice as much time, the partner agrees to reward him with 66% of the total resource at the evolutionary equilibrium, when partner choice is not costly. Conversely, when decision makers invest half as much time as their partner, they accept rewards of 33% only, showing that the fitness-maximizing strategy in this situation is to accept rewards proportional to each partner’s relative time investment.

**Figure 4:**
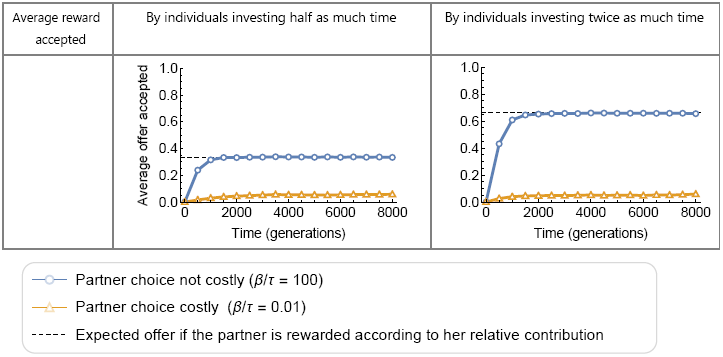
Evolution of the average reward accepted, depending on whether partners invest twice as much or half as much time into cooperation. Individuals investing twice as much time receive twice as much resources at equilibrium, and vice-versa.

#### A.2 Functioning of the neural networks

Each neuron in the networks computes an output signal of value

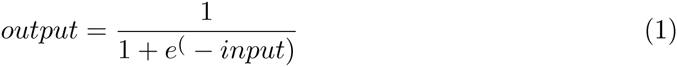

with *input* being a linear combination of the outputs of the neurons of the previous layer and the related synaptic weights. This is a function routinely used in evolutionary robotics (Nolfi, and Floreano, 2000). Synaptic weights can take values from the interval [−5, 5], and are randomly drawn from a uniform law covering this interval at the start of the simulation.

When applying mutations, to avoid networks to fall in suboptimal local maxima, mutations on the synaptic weights are drawn from a uniform distribution with a small probability 0.05; otherwise they ar drawn in a normal distribution centered around the synaptic weight’s value.

### B Analytical model.

We developed an analytical model to model the situation where individuals differ by their productivity (but not effort), and where only two productivities coexist in the population. The analytical model incorporates all of the features of the simulations, but with one simplification: we assume that the total number of interactions accepted per unit of time is the same for each individual. With this assumption, rejecting an opportunity to cooperate does not compromise the chances of cooperating later, but on the contrary grants new opportunities. This situation is analogous to the condition where 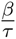 tends towards infinity in the simulations: social opportunities are plentiful at the scale of the length of interactions. When individuals reject an interaction, however, they are forced to postpone their social interaction to a later encounter. We assume that this entails an explicit cost expressed as a discounting factor *δ* (0 ≤ *δ* < 1). If we call the average payoff of an individual of productivity *i G*_*i*_ then *δG*_*i*_ will be the average expected payoff in the next interaction after rejecting an offer. When *δ* equals 1, refusing an interaction carries no cost; when *δ* equals 0, refusing an interaction will result in zero payoff from the next interaction. In practice, we will neglect the case where *δ* equals 1, as it leads to artefactual results (see below).

The assumption that only partners can decide of the division in our model is necessary so that the evolution of fairness is not explained trivially. When only one individual can decide, natural selection favors selfishness (André and Baumard, 2011). This is easy to understand. On the one hand, whatever reward a partner suggests, accepting it brings a greater gain than rejecting it for the decision maker. Therefore, in all cases, natural selection favors indiscriminate partners, with decision makers taking whatever benefits are made available to them. On the other hand, and as a result, selection favors stingy partners, offering the minimal possible amount. Because decision makers are in such an inferior bargaining position, in the following analysis we will focus on decision makers’— and not partners’—payoffs. A decision maker receiving a large share of the resource is a strong indication that there are evolutionary forces at work against the expected partners’ selfishness.

All our analyses assume that (i) individuals enter the population at a constant rate, (ii) evolution is slow compared to an individual’s lifespan (and thus) (iii) mutations are rare, and that (iv) there is no recombination between genetic traits *(p*_*ij*_ and *q*_*ij*_). As a consequence of (i) and (ii), the composition of the population does not change during an individual’s life. As a consequence of (iii) and (iv), at any evolutionary equilibrium, all the strategies present in the population must reach the same payoff for individuals of a given strength (only a high mutation rate or recombination rate could continuously re-introduce maladaptive strategies in the population, yielding a variance of payoffs at each generation).

Here we ask the same question answered in the main paper through simulations: how will the behavioural traits *r*_*ij*_ and MAR_*ij*_ *(i* and *j* ∈ *{HP, LP*}) evolve in an environment where LP and HP individuals coexist and share resources? As a reminder, MAR_*L P H P*_ reads as “the minimum reward that a LP individual will accept from a HP individual,” and *r*_*H P L P*_ as “the reward a HP individual will give to a LP individual.”

Following the precise evolutionary dynamics of the system to answer this question is quite a complex challenge, in particular due to epistasis phenomena. The low fitness benefits brought by a reward *r* can be compensated by high benefits from an acceptance threshold MAR, or small benefits obtained in interactions with individuals of one productivity could be compensated by high benefits received in interactions with the other productivity, generating linkage disequilibrium (McNamara et al., 2008). But as in (André and Baumard, 2011), it is easier to derive simple conditions on the payoff an individual would or would not have an interest in accepting at the evolutionary equilibrium.

#### B.1 Solving the system

The reasoning is more normative than descriptive, as we consider a situation in which the equilibrium has already been reached, and derive constraints on the values of traits that individuals should display at the equilibrium. To derive the payoff a LP individual should receive from a HP individual at the evolutionary equilibrium, we need to consider four arguments:

> **1. All individuals with the same productivity must gain the same payoff**. At the equilibrium, all HP individuals should gain the same payoff *G*_*HP*_ per interaction (otherwise it wouldn’t be an equilibrium), and the same is true for LP individuals. We thus only have two average payoffs in the population at the equilibrium. The average payoff of a HP individual is labeled *G*_*HP*_, and that of a LP individual is written *G*_*LP*_.
>
> 2. **Every individual of productivity** *i* **accepts exactly** δ*G*_*i*_, with *i* ∈ *{HP, LP*}. If an individual’s average payoff is *G*_*i*_, his expected payoff in the next interaction (if the current interaction is refused) will be *δG*_*i*_. As a consequence, a decision maker should never refuse a reward that is above the corresponding *δG*_*i*_, but should always refuse rewards that are below this level. At the equilibrium, because rewards from partners should evolve toward the minimum that decision makers will accept, individuals will always demand and accept exactly *δG*_*i*_, no matter who they are interacting with (regardless of their partner’s productivity). We thus have:
>
> 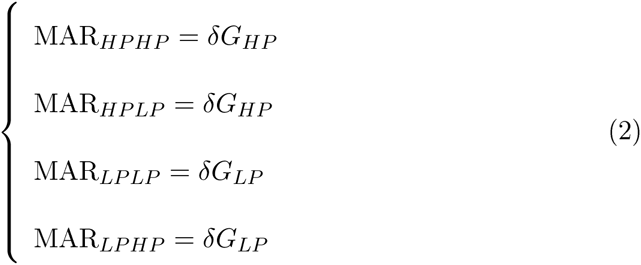
>
> 3. **Partners give their decision makers what they want at the evolutionary equilibrium, as long as** 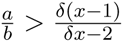
>
> Knowing (1) and (2), it can be shown that partners are always better off giving their decision makers what they “ask for” (*δG*_*i*_) at the evolutionary equilibrium, as long as *δ* < 1. The reasoning is as follows.
>
> Suppose that at the evolutionary equilibrium, all LP individuals refuse to give HP individuals what they ask for, namely *δG*_*HP*_(but all other demands are satisfied). The average social payoff of a LP individual in this population is then
>
> 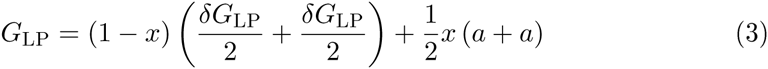
>
> with *x* the proportion of LP individuals in the population and *a* the productivity of LP individuals. *G*_*LP*_ can be decomposed into three terms: an average payoff obtained in interactions with other LP individuals ½(*a + a*), an average payoff obtained in interactions with HP individuals when HP individuals play the role of decision makers (in this case, under our hypothesis the reward will be rejected and the LP individual’s payoff will be discounted by *δ*), and, finally, an average payoff obtained in interactions with HP individuals when HP individuals are partners (the LP individual’s MAR is met, so they gain *δG*_*LP*_).
>
> Similarly, the payoff of a HP individual in this population is
>
> 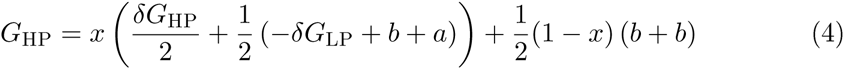
>
> with *b* the productivity of HP individuals. Solving the system composed of equations (3) and (4) gives us an expression for *G*_*HP*_ and *G*_*LP*_. The question we need to answer now is the following: what would happen if, in such a population, a mutant LP individual decided to accept to give HP individuals what they want? Upon meeting a HP individual and being assigned the role of partner, this mutant would gain *a* + *b* − *δG*_*HP*_ (the resource to be shared minus the demand of a HP individual) instead of just *δG*_*LP*_ (the average payoff being discounted). Knowing *G*_*LP*_ and *G*_*HP*_, it is easy to show that it is never possible that *δG*_*LP*_ ≥ *a* + *b* − *δG*_*HP*_ as long as *δ* < 1. In other words, at the evolutionary equilibrium, it is impossible that all LP individuals refuse to offer *δG*_*HP*_ to HP individuals, because they would gain more from doing so.
>
> What if there was some polymorphism in the population such that only *some* LP individuals refuse to give HP individuals what they ask for? The average social payoff of those LP individuals is still written the same as in equation (3). But because we know that at the evolutionary equilibrium all individuals with the same productivity must gain the same payoff, the payoff of all LP individuals will be the same, regardless of phenotype. The coexistence of two types of LP individuals in the population would imply that *δG*_*LP*_ = *a* + *a* − *δG*_*HP*_ (the payoff of the two types of LP individuals in the position of partner when paired with HP individuals is equal), but as we showed above, this is not possible as long as *δ* < 1. As a consequence, it is not only impossible that *all* LP individuals refuse to give HP individuals what they want at the evolutionary equilibrium, it is also impossible that *some* LP individuals refuse to give HP individuals what they want as long as *δ* < 1.
>
> Following the same reasoning, it can be shown that it is not possible for some individuals (of any productivity) to refuse to give their social partner (of any productivity) what they ask for at the evolutionary equilibrium as long as 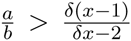 (^see^ SM section B.2). When 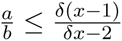, it is possible that LP individuals refuse to give other LP individuals what they ask for. This condition reflects the fact that if the difference of productivity between HP and LP individuals is too large, it is more beneficial for LP individuals to interact with HP individuals than with LP individuals. As we will see though, this is only possible when partner choice is costly. Moreover, as long as 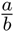 > 0.5, as is the case in our simulations, it is not worth it for LP individuals to refuse to interact with other LP individuals, and so all partners will give their decision makers what they want at the evolutionary equilibrium.
>
> If 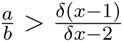, we can thus write:
>
> 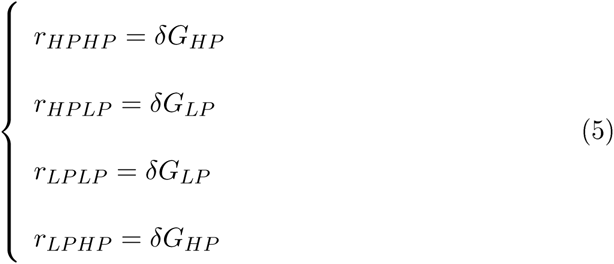
>
> and if 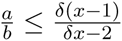, we can thus write:
>
> 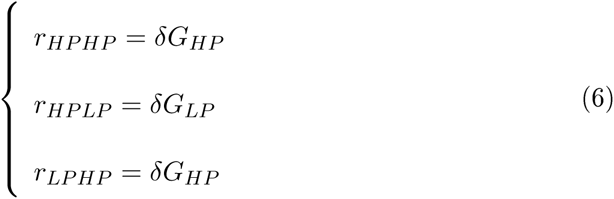
>
> 4. 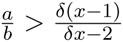, **no offer is never refused**
>
> If 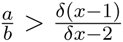, from step 3. it directly results that no reward is ever rejected at the evolutionary equilibrium, because each partner’s reward is exactly equal to the decision maker’s MAR, and thus each reward is accepted. If no reward is ever refused, the average payoff of LP and HP individuals respectively can be written as:
>
> 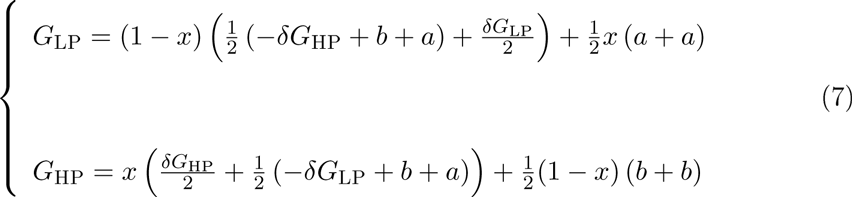
>
> Solving this system gives us an expression for *G*_*HP*_ and *G*_*LP*_ as a function of *x* and *δ* at the evolutionary equilibrium:
>
> 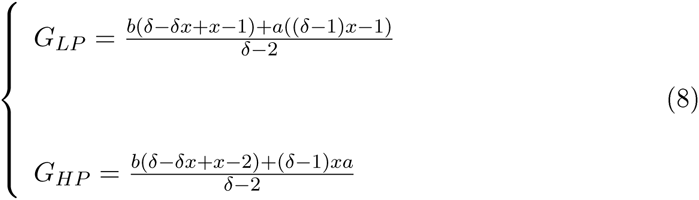
>
> From (5) and (8), it is straightforward to show that when *δ* tends toward 1 (partner choice is not costly), *r*_*LPHP*_ tends toward *b*. That is, when partner choice is not costly, even if LP individuals are in the strategically dominant position of partner, at the evolutionary equilibrium they offer HP individuals an amount that is exactly equal to their productivity *b*. In percentage, this corresponds to an offer proportional to the relative contribution of each individual: LP individuals offer HP individuals 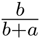 * 100 % of the total resource to be shared.
>
> Similarly, it can be shown that when *5* tends toward 1, LP individuals offer other LP individuals *a* resources, HP individuals offer other HP individuals *b* resources, and HP individuals offer LP individuals *a* resources. At the equilibrium, when partner choice is not costly each individual is rewarded with an amount exactly equal to his contribution.
>
> 5. 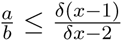, **all LP individuals refuse to interact with other LP individuals**
>
> In this case, the average payoff of LP and HP individuals respectively can be written as:
>
> 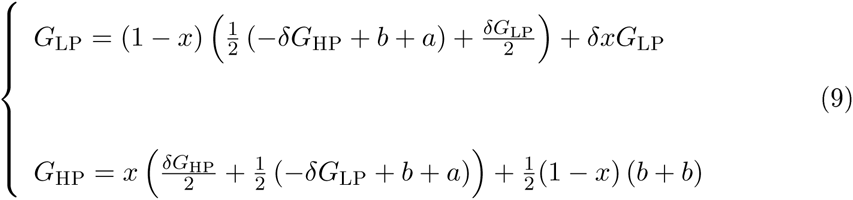
>
> Solving this system gives us an expression for *G*_*HP*_ and *G*_*LP*_ as a function of *x* and *5* at the evolutionary equilibrium:
>
> 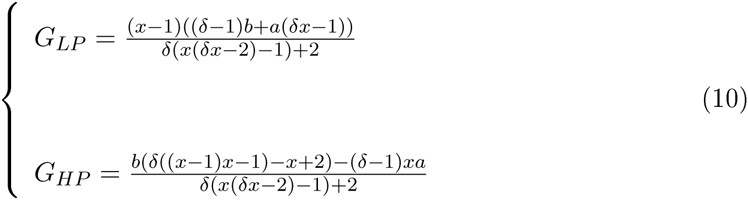
>
> From (6) and (10), it is straightforward to show that when *5* tends toward 1, the previous results hold: LP individuals offer HP individuals *b* resources, HP individuals offer other HP individuals *b* resources, and HP individuals offer LP individuals *a* resources.

#### B.2 Verification that partners are always better off giving their decision maker what they want at the evolutionary equilibrium, except when 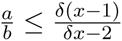

There are four hypothetical primary situations that need to be taken into account:

- A: when HP individuals are partners, they refuse to give other HP individuals what they want
- B: when HP individuals are partners, they refuse to give other LP individuals what they want
- C: when LP individuals are partners, they refuse to give other LP individuals what they want
- D: when LP individuals are partners, they refuse to give HP individuals what they want

These situations are not mutually exclusive, however, so the total number of possible situations is:

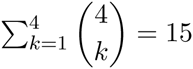

Situation D was already proven to be impossible at the evolutionary equilibrium in the previous section. We now show that the same holds for the 14 remaining situations, except in situation C. We give the expected social payoff of HP and LP individuals in each situation. We also give the condition that must be satisfied for each situation to be possible at the evolutionary equilibrium; it is then straightforward to show that, given our parameter values (0≤*x*≤1,0≤*δ*<1), this condition can never be satisfied.

Situation A:

- 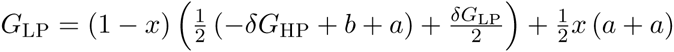
- 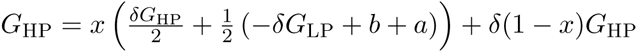
- Condition −*δG*_*HP*_ + *b* + *b* ≤ *δG*_*HP*_ impossible

Situation C:

- 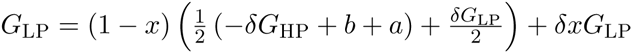
- 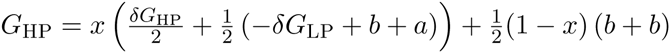
- Condition −*δG*_*LP*_ + *a* + *a* ≤ *δG*_*LP*_ impossible when 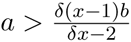

Situation B:

- 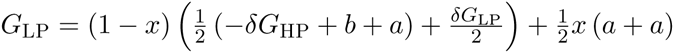
- 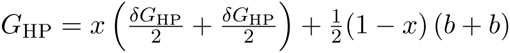
- Condition −*δG*_*LP*_ + *b* + *a* ≤ *δG*_*HP*_ impossible

Situation A & C:

- 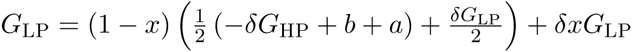
- 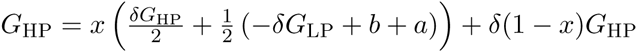
- Condition −*δG*_*LP*_ + *a* + *a* ≤ *δG*_*LP*_ ∧ *−δG*_*HP*_ + *b* + *b* ≤ *δG*_*HP*_ impossible

Situation B & C:

- 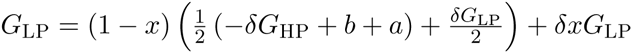
- 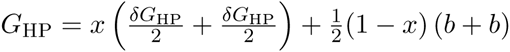
- Condition −*δG*_*LP*_ + *a* + *a* ≤ *δG*_*LP*_ ∧ −*δG*_*LP*_ + *b* + *a* ≤ *δG*_*HP*_ impossible

Situation C & D:

- 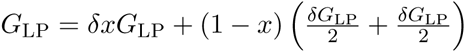
- 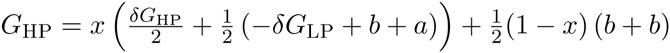
- Condition −*δG*_*LP*_ + *a* + *a* ≤ *δG*_*LP*_ ∧ − *δG*_*HP*_ + *b* + *a* ≤ *δG*_*LP*_ impossible

Situation B & D:

- 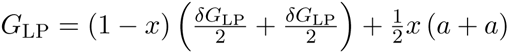
- 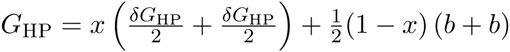
- Condition −*δG*_*HP*_ + *b* + *a* ≤ *δG*_*LP*_ ∧ −*δG*_*LP*_ + *b* + *a* ≤ *δG*_*HP*_ impossible

Situation A & D:

- 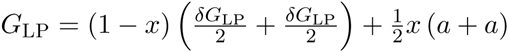
- 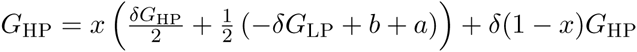
- Condition −*δG*_*HP*_ + *b* + *a* ≤ *δG*_*LP*_ ∧ −*δG*_*HP*_ + *b* + *b* ≤ *δG*_*HP*_ impossible

Situation A & B:

- 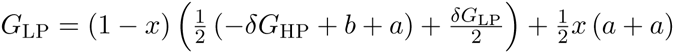
- 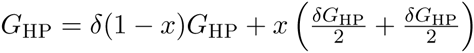
- Condition −*δG*_*HP*_ + *b* + *b* ≤ *δG*_*HP*_ ∧ −*δG*_*LP*_ + *b* + *a* ≤ *δG*_*HP*_ impossible

Situation A & C & D:

- 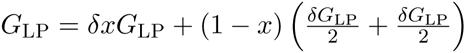
- 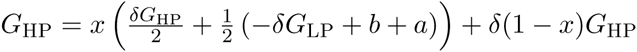
- Condition −*δG*_*LP*_ + *a* + *a* ≤ *δG*_*LP*_ ∧ *δG*_*HP*_ + *b* + *b* ≤ *δG*_*HP*_ ∧ *δG*_*HP*_ + *b* + *a* ≤ *δG*_*P*_ impossible

Situation A & B & C:

- 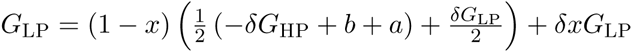
- 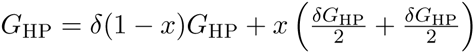
- Condition −*δG*_*LP*_ + *a* + *a* ≤ *δG*_*LP*_ ∧ −*δG*_*HP*_ + *b* + *b* ≤ *δG*_*HP*_ ∧ −*δG*_*LP*_ + *b* + *a* ≤ *δG*_*HP*_ impossible

Situation B & C & D:

- 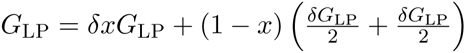
- 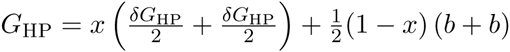
- Condition −*δG*_*LP*_ + *a* + *a* ≤ *δG*_*LP*_ ∧ −*δG*^*HP*^ + *b* + *a* ≤ *δG*_*LP*_ ∧ −*δG*_*LP*_ + *b* + *a* ≤ *δG*_*HP*_ impossible

Situation A & B & D:

- 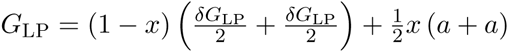
- 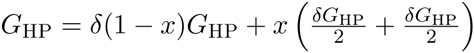
- Condition −*δG*_*HP*_ + *b* + *b* ≤ *δG*_*HP*_ ∧ *δG*_*LP*_ + *b* + *a* ≤ *δG*_*HP*_ ∧ *δG*_*HP*_ + *b* + *a* ≤ *δG*_*LP*_ impossible

Situation A & B & C & D:

- 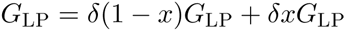
- 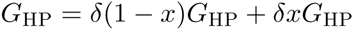
- Condition −*δG*_*HP*_ + *b* + *b* ≤ *δG*_*HP*_ ∧ −*δG*_*LP*_ + *b* + *a* ≤ *δG*_*HP*_ ∧ −*δG*_*LP*_ *+ a* + *a* ≤ *δG*_*LP*_ ∧ −*δG*_*HP*_ + *b* + *a* ≤ *δG*_*LP*_ impossible

As explained in the previous section, the verification that it is not possible for *some* (but not all) individuals not to interact with other individuals at the evolutionary equilibrium (in case of polymorphism) is already implied by the use of not strict inequalities.

### C Supplementary discussion

#### C.1 Opportunity costs

In the main article, we explain that when high-productivity individuals are assessing a low-productivity individual’s reward, they have opportunity costs (or “outside options”) of 2 because they expect to receive 2 with other high-productivity individuals *on average*. It is important to see that this is true only because high-productivity individuals have an equal chance of playing the role of either decision-maker or partner when they interact with other high-productivity individuals. If some high-productivity individuals always played the role of decision maker with other high-productivity individuals, they would be exploited all the time by those high-productivity partners, which would drastically reduce their outside options when bargaining with low-productivity individuals, preventing the evolution of proportionality. Thus, in our model the evolution of proportionality depends as much on the possibility of changing *roles* as on the possibility of changing *partners*. In real life, this is the equivalent of having a rich and varied social life with multiple cooperative opportunities in which one is not always in the worse bargaining position (Wiessner, 1996; Kaplan et al., 2009).

#### C.2 Theoretical problems with partner choice

Partner choice is an intrinsically complicated subject. The existence of a wide variety of cooperative partners to choose from means that a wide variety of social strategies can coexist and provide the same benefits, complicating evolutionary analysis. For example, an individual’s acceptance of low rewards as a decision maker could be compensated by the low rewards she herself makes as a partner. Or some low payoffs received when interacting with low-productivity individuals could be compensated by high payoffs received when interacting with high-productivity individuals.

These effects explain why a quick look at the evolved strategies of individuals is not always enough to find a pattern of proportionality. This is especially true with neural networks working on a continuum of productivities or effort. While, as we have shown, the theoretical fitness-maximizing behavior is to offer an amount proportional to one’s own relative contribution, it is not necessarily the case that neural networks will produce proportional offers for the *whole* range of inputs they are exposed to. Imagine an individual who offers proportional rewards only to the best producers in the population, while offering less-than-proportional rewards to other individuals. At the evolutionary equilibrium, our model predicts that these unfair rewards will be rejected. But as long as finding a new partner is not costly, being rejected does not lead to a loss of fitness. As a consequence, any individual can offer less-than-proportional rewards to a fraction of the population, as long as another fraction still accepts the rewards she makes that are proportional. In other words, individuals can specialize in offering proportional rewards to only a fraction of the range of productivities in the population, and stop interacting with the remaining fraction. Because they stop interacting, the rewards offered to this fraction become subject to drift.

Because of this mechanism, it is possible that averaging the output of different evolved neural networks does not reveal a pattern of proportionality. In our simulations, averaging the output of 15,000 neural networks producing MARs yielded an almost perfect proportional relationship between contributions and MARs (main paper, Fig. 3C). Plotting the average output of 15,000 neural networks producing *rewards* did not show such a perfectly proportional relationship, although it was not far from it. Here, it is important to remember that despite this variability in the rewards that are extended, proportionality prevails when we look only at the interactions that *actually take place:* only proportional rewards are accepted at the evolutionary equilibrium, as evidenced in Fig. 3B of the main article.

Finally, problems of neutrality add complexity to the analysis. Although at the beginning of our simulations raising MARs drove the evolution of proportional rewards, once proportional rewards had spread in the population, the selection pressure to maintain high MARs disappeared: if all individuals offer rewards of *r*, requesting *r* or *r – e* as a decision maker brings the same payoff. Because of drift, MARs can thus start to decrease, and in turn partners will be selected to decrease their rewards to try to exploit those undemanding decision makers. This exploitation cannot last for long, as it soon revives the selection pressure to increase MARs, but the dynamic exists. Although it is rather easy to conceptualize why, under appropriate conditions, partner choice leads to proportionality, the actual dynamics underlying this result are far from straightforward to understand.

## D Acknowledgments

SD thanks the *Région Ile-de-France* for funding this research through a *2012 DIM “Prob-lématiques transversales aux systémes complexes”* grant, and thanks the *Institut des systémes complexes* and the *Ecole Doctorale Frontières du Vivant (FdV) - Programme Bet-tencourt* for its support. This work was supported by ANR-10-LABX-0087 IEC and ANR-10-IDEX-0001-02 PSL*.

## E Author contributions

SD built the simulations and analytical model, analyzed the data and wrote the manuscript. J-B.A and NB designed and coordinated the study.

## F Data availability

The data presented in this paper will be archived online on dryad.com and on the first author’s website.

## G Competing interests

We have no competing interests to declare.

## References

Adams, J. S., 1963. Toward an Understanding of Inequity. Journal of abnormal psychology 67:422–436.

Adams, J. S. and P. R. Jacobsen, 1964. Effects of Wage Inequities on Work Quality. Journal of abnormal psychology 69:19–25.

Aktipis, C. A., 2004. Know when to walk away: contingent movement and the evolution of cooperation. Journal of theoretical biology 231:249–60.

Aktipis, C. A., 2011. Is cooperation viable in mobile organisms? Simple Walk Away rule favors the evolution of cooperation in groups. Evolution and Human Behavior 32:263–276.

Alvard, M. S., 2002. Carcass ownership and meat distribution by big-game cooperative hunters, vol. 21.

Amici, F., E. Visalberghi, and J. Call, 2014. Lack of prosociality in great apes, capuchin monkeys and spider monkeys: convergent evidence from two different food distribution tasks. Proc. R. Soc.B.

André, J.-B. and N. Baumard, 2011. Social opportunities and the evolution of fairness. Journal of theoretical biology 289:128–35.

Aristotle, 1999. Nicomachean Ethics, vol. 112.

Austin, W. and E. Walster, 1974. Reactions to confirmations and disconfirmations of expectancies of equity and inequity.

Barclay, P., 2004. Trustworthiness and competitive altruism can also solve the “tragedy of the commons”. Evolution and Human Behavior 25:209–220.

Barclay, P., 2006. Reputational benefits for altruistic punishment. Evolution and Human Behavior 27:325–344.

Barclay, P., 2011. Competitive helping increases with the size of biological markets and invades defection. Journal of theoretical biology 281:47–55.

Barclay, P., 2013. Strategies for cooperation in biological markets, especially for humans. Evolution and Human Behavior.

Barclay, P. and B. Stoller, 2014. Local competition sparks concerns for fairness in the ultimatum game. Biology letters 10:1–4.

Barclay, P. and R. Willer, 2007. Partner choice creates competitive altruism in humans. Proceedings. Biological sciences / The Royal Society 274:749–53.

Barkow, J. H., L. Cosmides, and J. Tooby, 1992. The adapted mind: evolutionary psychology and the generation of culture. Oxford University Press.

Baumard, N., J. André, and D. Sperber, 2013. A mutualistic approach to morality: The evolution of fairness by partner choice. Behavioral and Brain Sciences 6:59–122.

Baumard, N. and P. Boyer, 2013. Explaining moral religions. Trends in cognitive sciences 17:272–80.

Baumard, N., O. Mascaro, and C. Chevallier, 2012. Preschoolers are able to take merit into account when distributing goods. Developmental psychology 48:492–498.

Binmore, K., 2005. Natural Justice. Oxford University Press.

Bird, R. B. and E. a. Power, 2015. Prosocial signaling and cooperation among Martu hunters. Evolution and Human Behavior.

Bräuer, J. and D. Hanus, 2012. Fairness in Non-human Primates? Social Justice Research 25:256–276.

Camerer, C, 2003. Behavioral game theory: Experiments in strategic interaction, vol. 32. Princeton University Press, Princeton, New Jersey.

Cappelen, A., A. Hole, E. Sørensen, and B. Tungodden, 2007. The pluralism of fairness ideals: An experimental approach. American Economic Review 97:818–827.

Cappelen, A., E. Sørensen, and B. Tungodden, 2010. Responsibility for what? Fairness and individual responsibility. European Economic Review 54:429–441.

Cohen, C, 2009. Why not socialism? Princeton University Press.

Dawes, C. T., J. H. Fowler, T. Johnson, R. Mcelreath, and O. Smirnov, 2007. Egalitarian motives in humans. Nature 446:794–796.

Debove, S., J.-b. André, and N. Baumard, 2015a. Partner choice creates fairness in humans. Proc. R. Soc.B 282.

Debove, S., N. Baumard, and J.-B. André, 2015b. Evolution of equal division among unequal partners. Evolution 69:561–569.

Fehr, E. and K. Schmidt, 1999. A theory of fairness, competition, and cooperation. The quarterly journal of economics 114:817–868.

Fiske, A. P., 1992. The four elementary forms of sociality: framework for a unified theory of social relations. Psychological review 99:689–723.

Forber, P. and R. Smead, 2014. The evolution of fairness through spite. Proceedings of the Royal Society B 281.

Frohlich, N., J. Oppenheimer, and A. Kurki, 2004. Modeling other-regarding preferences and an experimental test. Public Choice 119:91–117.

Gale, J., K. Binmore, and L. Samuelson, 1995. Learning to be imperfect: The ultimatum game. Games and Economic Behavior 8:56–90.

Gavrilets, S. and S. M. Scheiner, 1993. The genetics of phenotypic of reaction norm shape V. Evolution of reaction norm shape. Journal of evolutionary biology 48:31–48.

Geraci, A. and L. Surian, 2011. The developmental roots of fairness: Infants’ reactions to equal and unequal distributions of resources. Developmental Science 14:1012–1020.

Gurven, M., 2004. To give and to give not: The behavioral ecology of human food transfers. Behavioral and Brain Sciences 27:543–583.

Güth, W., R. Schmittberger, and B. Schwarze, 1982. An experimental analysis of ultimatum bargaining. Journal of Economic Behavior & Organization 3:367–388.

Hoebel, E. A., 1954. The Law of Primitive Man: A Study in Comparative Legal Dynamics P. 372.

Hoel, M., 1987. Bargaining games with a random sequence of who makes the offers. Economics Letters 24:5–9.

Homans, G. C., 1958. Social Behavior as Exchange. The american journal of sociology 63:597–606.

Huck, S. and J. Oechssler, 1999. The Indirect Evolutionary Approach to Explaining Fair Allocations. Games and Economic Behavior 28:13–24.

Kanngiesser, P., N. Gjersoe, and B. M. Hood, 2010. The effect of creative labor on property-ownership transfer by preschool children and adults. Psychological science 21:1236–41.

Kaplan, H. and M. Gurven, 2005. The Natural History of Human Food Sharing and Cooperation: A Review and a New Multi-Individual Approach to the Negotiation of Norms. Pp. 75–113, in R. B. &. E. F. H. Gintis, S. Bowles, ed. Moral sentiments and material interests: The foundations of cooperation in economic life. MIT Press.

Kaplan, H. S., P. L. Hooper, and M. Gurven, 2009. The evolutionary and ecological roots of human social organization. Philosophical transactions of the Royal Society B. 364:3289–99.

Killingback, T. and E. Studer, 2001. Spatial Ultimatum Games, collaborations and the evolution of fairness. Proceedings. Biological sciences / The Royal Society 268:1797–801.

Konow, J., 2003. Which is the fairest one of all? A positive analysis of justice theories. Journal of economic literature XLI:1188–1239.

Liénard, P., C. Chevallier, O. Mascaro, P. Kiura, and N. Baumard, 2013. Early understanding of merit in Turkana children. Journal of Cognition and Culture 13:57–66.

Lyle, H. F. and E. a. Smith, 2014. The reputational and social network benefits of prosociality in an Andean community. Proceedings of the National Academy of Sciences of the United States of America 111:4820–5.

Marshall, G., A. Swift, D. Routh, and C. Burgoyne, 1999. What is and what ought to be popular beliefs about distributive justice in thirteen countries. European Sociological Review 15:349–367.

McNamara, J., Z. Barta, L. Fromhage, and A. Houston, 2008. The coevolution of choosiness and cooperation. Nature 451:189–192.

Mellers, B. a., 1982. Equity judgment: A revision of Aristotelian views. Journal of Experimental Psychology: General 111:242–270.

Nesse, R. M., 2007. Runaway social selection for displays of partner value and altruism. Biological Theory 2.

Nettle, D., K. Panchanathan, T. S. Rai, and A. P. Fiske, 2011. The Evolution of Giving, Sharing, and Lotteries. Current Anthropology 52:747–756.

Noë, R. and P. Hammerstein, 1994. Biological markets: supply and demand determine the effect of partner choice in cooperation, mutualism and mating. Behavioral ecology and sociobiology 35:1–11.

Noë, R., J. V. Hooff, and P. Hammerstein, 2001. Economics in nature: social dilemmas, mate choice and biological markets. Cambridge University Press, Cambridge, England.

Nolfi, S. and D. Floreano, 2000. Evolutionary robotics: the biology, intelligence, and technology of self-organizing machines. The MIT Press.

Nowak, M. a., K. M. Page, and K. Sigmund, 2000. Fairness versus reason in the ultimatum game. Science 289:1773–5.

Osborne, M. and A. Rubinstein, 1990. Bargaining and markets. Academic Press, Inc, San Diego, California.

Page, K. M. and M. a. Nowak, 2002. Empathy leads to fairness. Bulletin of mathematical biology 64:1101–16.

Page, K. M., M. a. Nowak, and K. Sigmund, 2000. The spatial ultimatum game. Proceedings. Biological sciences / The Royal Society 267:2177–82.

Rand, D. G., C. E. Tarnita, H. Ohtsuki, and M. a. Nowak, 2013. Evolution of fairness in the one-shot anonymous Ultimatum Game. Proceedings of the National Academy of Sciences of the United States of America 110:2581–6.

Roberts, G., 1998. Competitive altruism: from reciprocity to the handicap principle. Proceedings of the Royal Society B: Biological Sciences 265:427–431.

Robinson, P. H. and R. Kurzban, 2007. Concordance and Conflict in Intuitions of Justice. Minn. L. Rev. 91:1–75.

Rubinstein, A., 1982. Perfect equilibrium in a bargaining model. Econometrica: Journal of the Econometric Society 50:97–110.

Sahlins, M., 1972. Stone age economics. Aldine - Atherton, Inc, Chicago and New York.

Schäfer, M., D. B. M. Haun, and M. Tomasello, 2015. Fair Is Not Fair Everywhere.

Schmidt, M. F. H. and J. a. Sommerville, 2011. Fairness expectations and altruistic sharing in 15-month-old human infants. PLoS ONE 6.

Skitka, L. J., 2012. Cross-Disciplinary Conversations: A Psychological Perspective on Justice Research with Non-human Animals. Social Justice Research 25:327–335.

Sloane, S., R. Baillargeon, and D. Premack, 2012. Do Infants Have a Sense of Fairness? Psychological Science 23:196–204.

Sylwester, K. and G. Roberts, 2013. Reputation-based partner choice is an effective alternative to indirect reciprocity in solving social dilemmas. Evolution and Human Behavior 34:201–206.

Trivers, R., 1971. The evolution of reciprocal altruism. Quarterly review of biology 46:35–57.

Trivers, R., 2006. Reciprocal altruism: 30 years later, in Cooperation in Primates and Humans: Mechanisms and Evolution. Springer.

Turiel, E., 2002. The Culture of morality: Social development, context, and conflict.

Walster, E., E. Berscheid, and G. W. Walster, 1973. New Directions in Equity Research. Advances in Experimental Social Psychology 25:151–176.

Wiessner, P., 1996. Leveling the hunter: constraints on the status quest in foraging societies. Pp. 171–192, in Food and the Status Quest: An Interdisciplinary Perspective, p. wiessne ed. Berghahn Books, Oxford, UK.

